## Supplemental Data for "Glycolytic flux controls retinal progenitor cell differentiation via regulating Wnt signaling"

### Supplemental Figures

**Figure 1– Figure supplement 1. Expression of *Pten* and mature photoreceptor markers during murine retinal development.**

UMAP plot of scRNA-seq data collected from wild-type retinas between E11 and P14 (Clark et al., 2019). Transcript distribution of *Pten*, and mature photoreceptor markers (*Prph2, Impg2, Pde6g, Gm11744, Gnat2*) at different stages of retinal development.

**Figure 6 – Figure supplement 1. Upregulation of Wnt-associated genes in *Pten* cKO retinas.** PANTHER (protein analysis through evolutionary relationships) classification analysis of DEGs of P0 bulk RNAseq showing upregulation of Wnt signaling and glycolysis (arrows).

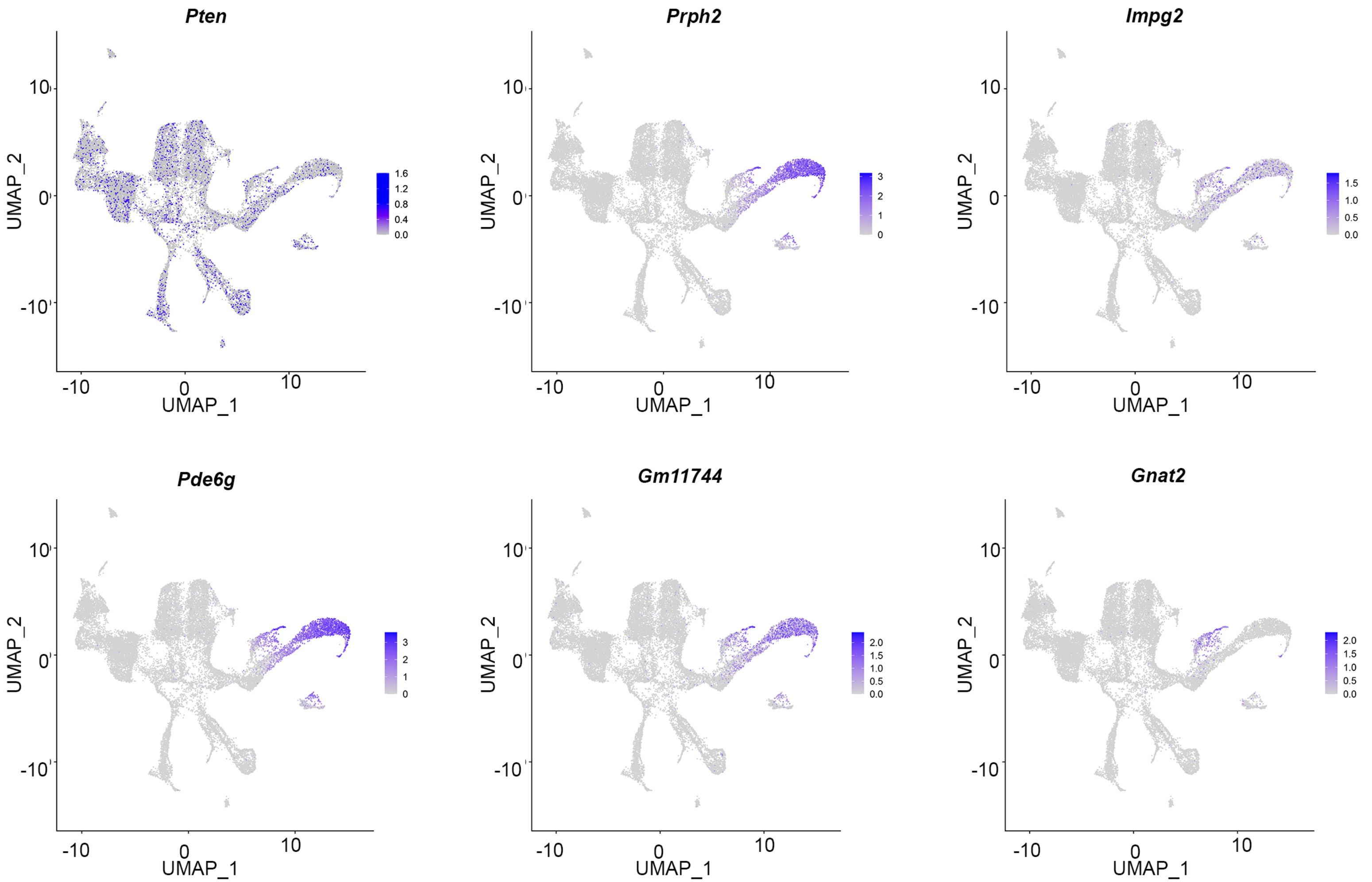


Figure 1-figure supplement 1

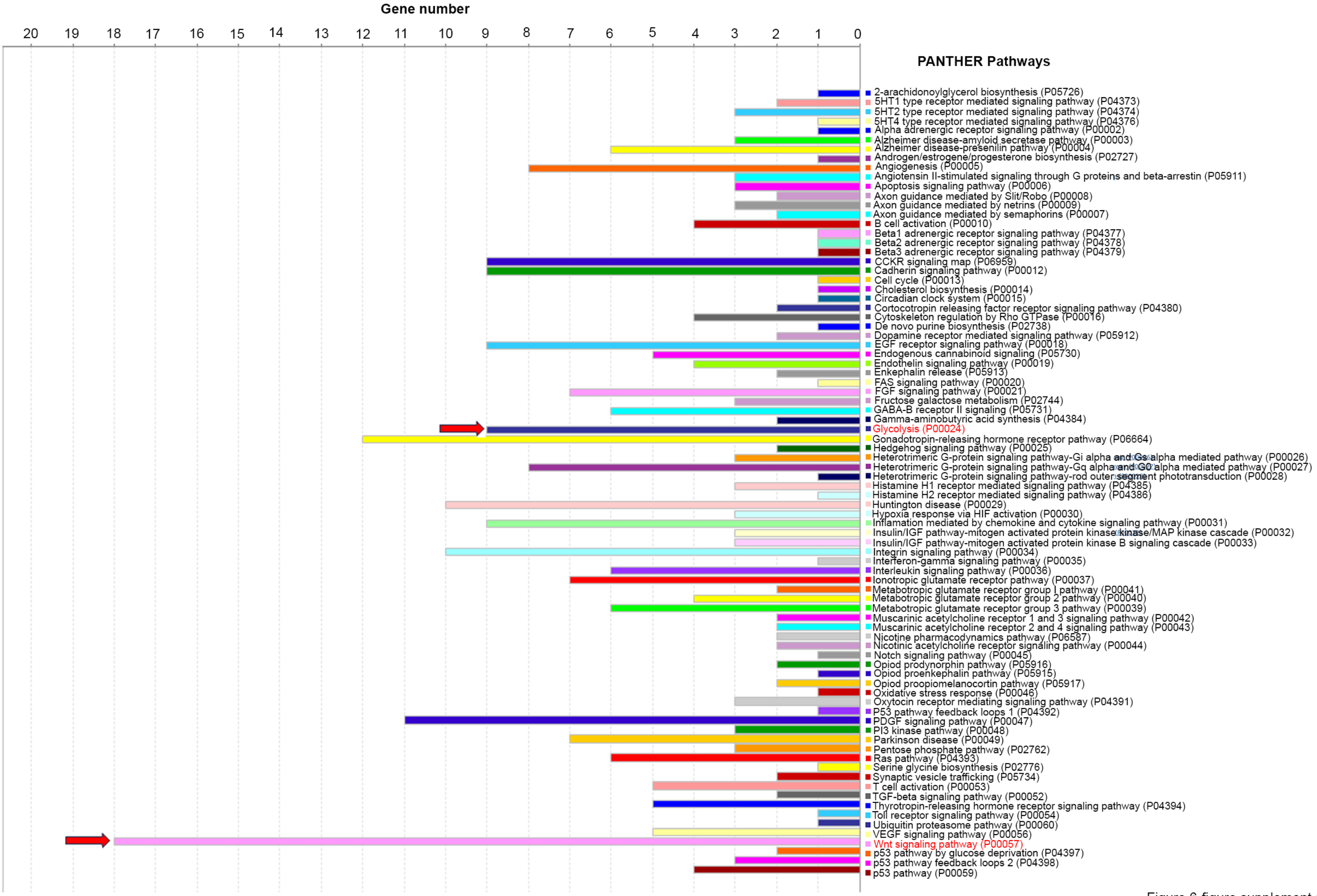


Figure 6-figure supplement 1
